# May the force be with you: the role of hyper-mechanostability of the bone sialoprotein binding protein during early stages of Staphylococci infections

**DOI:** 10.1101/2022.12.09.519785

**Authors:** Priscila S. F. C. Gomes, Meredith Forrester, Margaret Pace, Diego E. B. Gomes, Rafael C. Bernardi

## Abstract

The bone sialoprotein-binding protein (Bbp) is a mechanoactive MSCRAMM protein expressed on the surface of *Staphylococcus aureus* that mediates adherence of the bacterium to fibrinogen-*α* (Fg*α*), a component of the bone and dentine extracellular matrix of the host cell. Mechanoactive proteins like Bbp have key roles in several physiological and pathological processes. Particularly, the Bbp:Fg*α* interaction is important in the formation of biofilms, an important virulence factor of pathogenic bacteria. Here, we investigated the mechanostability of the Bbp:Fg*α* complex using *in silico* single-molecule force spectroscopy (SMFS), in an approach that combines results from all-atom and coarse-grained steered molecular dynamics (SMD) simulations. Our results show that Bbp is the most mechanostable MSCRAMM investigated thus far, reaching rupture forces beyond the 2 nN range in typical experimental SMFS pulling rates. Our results show that high force-loads, which are common during initial stages of bacterial infection, stabilize the interconnection between the protein’s amino acids, making the protein more “rigid”. Our results offer new insights that are crucial on the development of novel anti-adhesion strategies.

## Introduction

*Staphylococcus aureus* infections have a high clinical and communal impact with an estimated mortality rate that can reach 30.2%.^1^ The persistence of these infections lies on the *S. aureus*’ ability to form biofilms,^2–4^ and the eventual dissemination of these pathogenic bacteria throughout the body.^5^ Despite the increase in sterilization and hygienic measures, modern medical devices play a key role in the transfer of these bacterial colonies through device-associated biofilm infections.^6–8^ The contamination of patients during medical and dental procedures is of increasing relevance, particularly with the emergence of drug-resistant bacteria. In the dental field, it has been estimated that the carrier prevalence of *S. aureus* in healthy adults varies from 24% to 84%. ^9^ Additionally, the oral cavity is a source for cross infection and dissemination of the infection directly into the bloodstream, increasing the likelihood of septicemia and possibly death. ^10–12^

Biofilms shelter the bacteria and enhance the persistence of infection by eluding innate and adaptive host defenses.^13,14^ Biofilms also form a barrier, protecting colonies from biocides and antibiotic chemotherapies.^15^ Adhesins play critical roles during infection, especially during the early step of adhesion when bacterial cells are exposed to mechanical stress.^16^ Adhesins bind to their target ligands, holding it tight to them even at extreme force loadings that largely outperform classical binding forces. ^17^ The resilience to mechanical forces provides the pathogen with a means to withstand high levels of mechanical stress during biofilm formation, thus yielding these pathogens highly resistant to breaking these cell adhesion bonds. These unusual stress-dependent molecular interactions play an integral role during bacterial colonization and dissemination and when studied, reveal critical information about pathosis.^18^

Among *S. aureus* adhesins, the bone sialoprotein binding protein (Bbp) is a bifunctional Microbial Surface Component Recognizing Adhesive Matrix Molecule (MSCRAMM). ^19^ Bbp is part of the MSCRAMM serine-aspartate repeat (Sdr) family that also includes SdrF and SdrG in *S. epidermidis*, and clumping factor A (ClfA), B (ClfB), SdrC, and SdrE in *S. aureus*.^20–23^ Ligand-binding for Bbp occurs generally in the N-terminal region, from residues 273 to 598, where Bbp binds to fibrinogen-*α* (Fg*α*), a glycopeptide on bone and dentine extracellular matrix (ECM). Bbp’s binding region is subdivided into domains N2 and N3, which are made up of two layers of *β*-sheets with an open groove at the C-terminus where primary ligand binding occurs^24^ (Figure 1). The binding of Fg*α* follows a “dock, lock, and latch” mechanism,^25–29^ that has been previously investigated by a myriad of techniques.^30–34^ Thus, the pathogenic bacteria does not invade a host cell, but rather adheres to the ECM via Bbp:Fg*α* interactions.^35^

**Figure 1:**
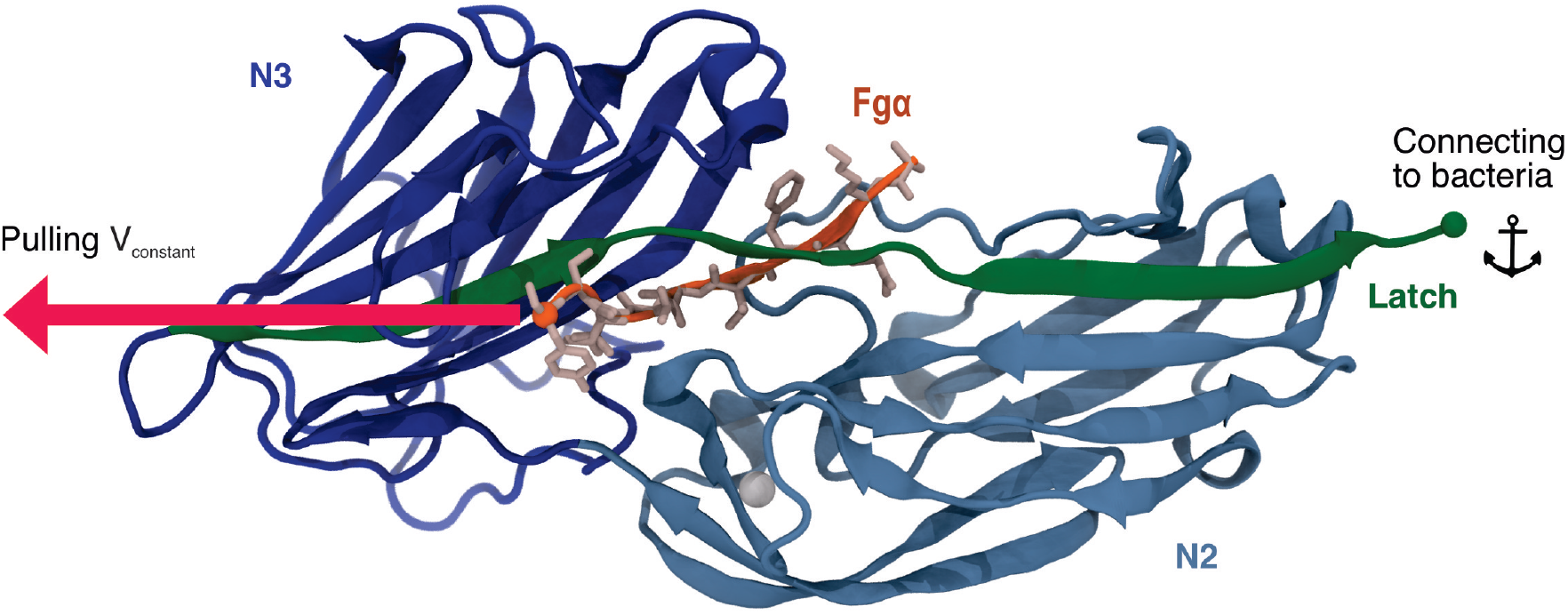
Tridimensional structure of Bbp’s adhesion domain. The protein is represented in cartoon, colored by its different domains. The latch is highlighted in green. Fg*α* is colored in orange and its aminoacids represented as sticks colored in light pink. The SMD pulling and anchor points are indicated in the image as spheres.

Using a combination of *in silico* and *in vitro* single-molecule force spectroscopy (SMFS), we have previously reported that *S. epidermidis*’ adhesin SdrG, when in complex with Fg*β*, was able to withstand extreme mechanical loads.^34^ The necessary force applied to rupture the SdrG:Fg*β* complex was shown to be an order of magnitude stronger than that needed to rupture the widely employed Streptavidin:biotin complex, ^36^ and more than twice of that of cellulosomal cohesin:dockerin interactions. ^37^ The force resilience is way superior to proteins found in the cytoplasm, which are typically 20 to 1000 times weaker than Staphylococci adhesins.^38,39^ A molecular mechanism for a catch-bond behavior of the SdrG:Fg*β* was then revealed by investigating the system in a “force-clamp” regime, ^40^ with magnetic tweezers based SMFS revealing that the SdrG:Fg*β* bond can live for hours under force loads.^41^ Here, taking advantage of a powerful *in silico* SMFS approach, we describe how Bbp plays a key role in bacterial adhesion during nosocomial infections, by investigating the Bbp:Fg*α* complex at different pulling velocities combining all-atom (aa) and coarse-grained (CG) steered molecular dynamics (SMD) simulations. Building on *in vitro* SMFS data, our results point to Bbp’s interaction with the extracellular matrix fibrinopeptide as the most mechanostable so far investigated, independent of the loading rate. Our findings reveal that a few key interactions are responsible for the outstanding force resilience of the complex. Furthermore, our results offer insights into the development of anti-adhesion strategies.

## Results

### Bbp is highly mechanostable under stress

To probe the mechanics of the interaction between Bbp and Fg*α*, and to characterize the atomic details of the complex under force load, we performed aa-SMD simulations with Bbp anchored by its C-terminal while Fg*α* was pulled at different velocities (Table 2). Molecular dynamics using classic mechanics force fields have been shown to be able to predict rupture forces,^37,40,42–44^ and also predict molecular events even in complex electrostatic conditions. ^45–50^ The simulations resulted in Force vs. extension curves that reveal a clear one-step rupture event, as represented in Figure 2A. For the slowest pulling velocity, 160 replicas were performed following a wide-sampling paradigm previously developed in our group. ^51^ At the pulling velocity of 2.5 *×* 10^−04^ nm/ps, we observed that the most probable rupture force for the complex was 3,510 pN, as described by the Bell-Evans (BE)^52,53^ fit of the peak forces at that pulling speed (see Figure 2B). Our results reveal that Bbp:Fg*α* is the most mechanostable complex investigated thus far, which is in agreement with preliminary experimental data. ^34^ To investigate the dependence of the mechanostability of Bbp:Fg*α* on the force loading rate, we performed CG-SMD simulations at several, much lower, pulling speeds (Table 2). We have recently shown that aa-SMD and CG-SMD can be combined to in an *in silico* SMFS approach.^54^ Here, the combination of the two levels of molecular details is capable of rendering predictions that are consistent with theory and experimentation. A Dudko-Hummer-Szabo ^55^ (DHS) fit was performed through the SMD data, including both the aa-SMD, and the CG-SMD (see Figure 3). The DHS fit suggests that the system should rupture at forces higher than 2 nN at 10^5^ pN/s force loading rate, in agreement with experimental data.^34^ It is interesting to note that the BE model is able to fit well all the simulation results, at both aa and CG level, as evidenced by the density plots in Figure 3.

**Figure 2:**
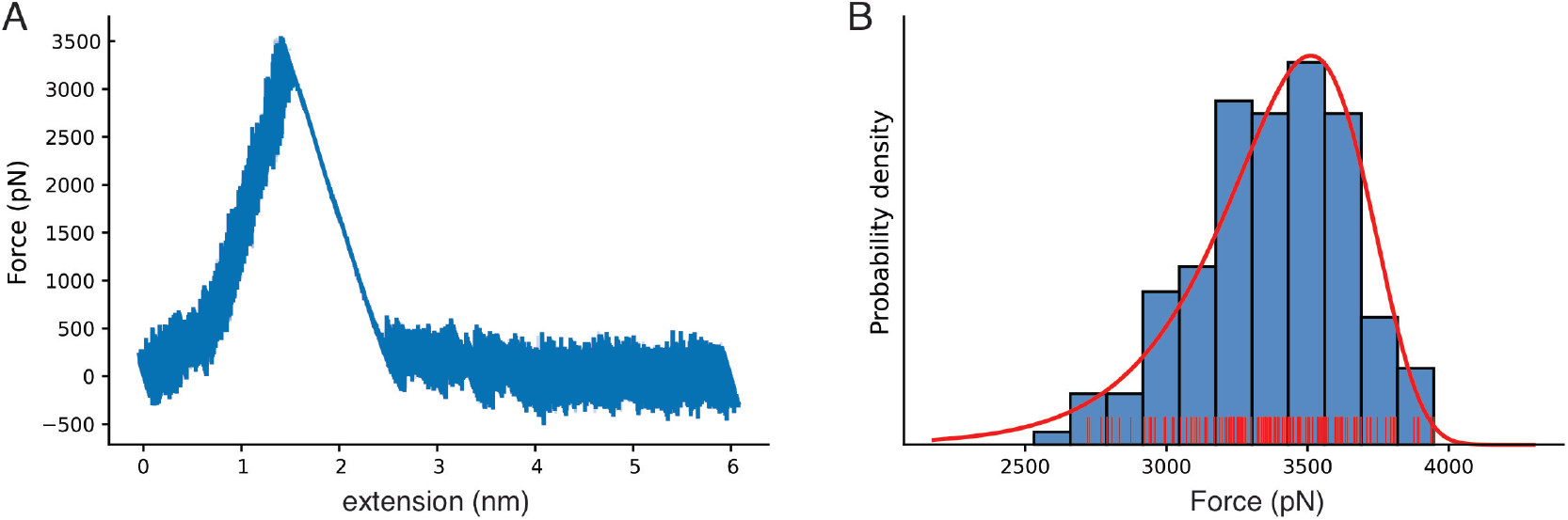
Bbp mechanostability under high mechanical load. **a)** Force versus extension curve as an exemplary trace, with rupture peak force at 3,510 pN. **b)** Histogram for the most probable rupture force (blue, rugged plot in red) with the Bell-Evans (BE) model for the first rupture peak (red), based on the aa-SMD simulation replicas with the slowest simulated pulling velocity (2.5 *×* 10^−4^ nm/ps).

**Figure 3:**
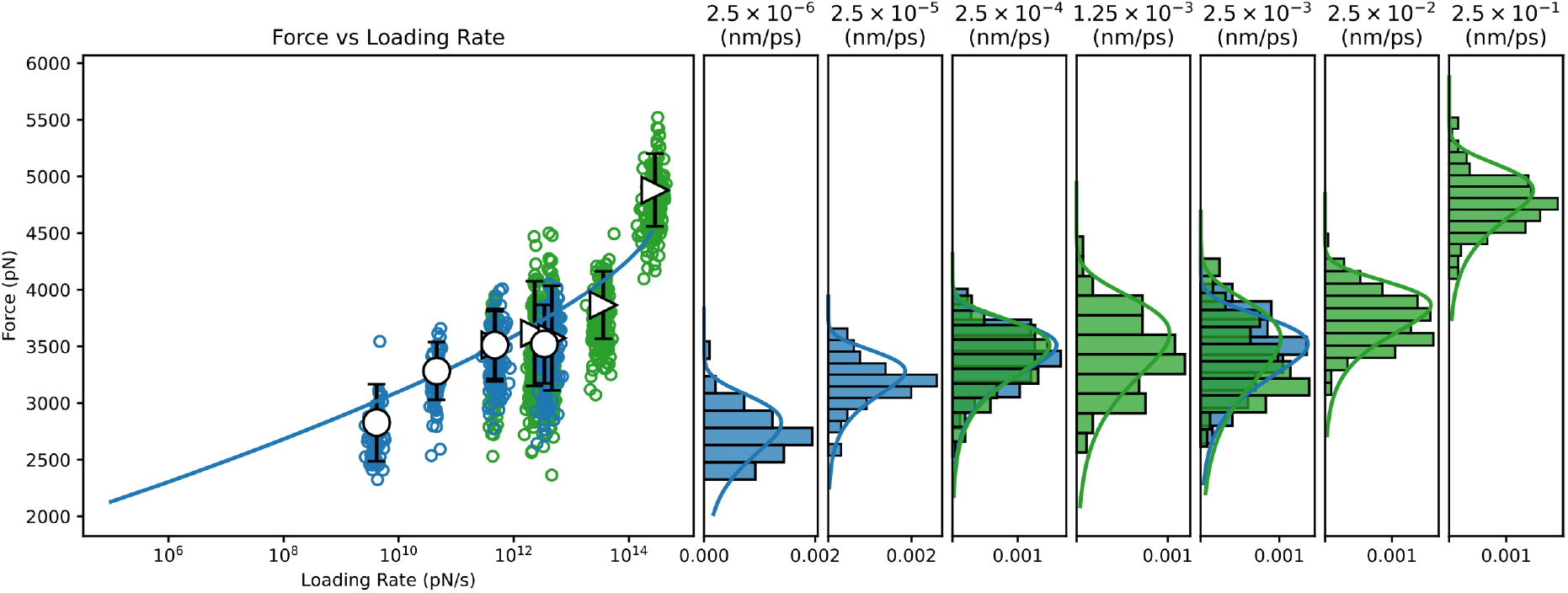
Dynamic Force spectrum for the Bbp:Fg*α* complex combining data from all-atom and coarse-grained SMD simulations. All-atom, and Coarse-grained steered molecular dynamics simulations (CG-SMD and aa-SMD) were performed at different velocities: 2.5 *×* 10^−6^ to 2.5 *×* 10^−3^ nm/ps (blue) and 2.5 *×* 10^−4^ to 2.5 *×* 10^−3^ nm/ps (green), respectively. A Dudko-Hummer-Szabo ^55^ (DHS) fit was performed through the SMD dataset predicting Δ*x* = 7.489 *×* 10^−2^ *nm, k*_*off*_^0^ = 2.596 *×* 10^−12^ *s*^−1^, Δ*G* = 2.293 *×* 10^2^ *kBT*.

### Key hydrogen bonds are responsible for Bbp:Fg*α* high mechanostability

After confirming that Bbp:Fg*α* complex presents a hyperstable interaction under shear mechanical load, we used the approximately 3 *μ*s of aa simulation data to investigate the molecular origin of the mechanostability of the complex. Previously, simulations of the SdrG:Fg*β* revealed the presence of frequent and persistent hydrogen bonds (H-bonds) between the peptide and the protein backbone, showing that the high-force resilience of the complex was largely independent of the peptide side-chains interactions, and therefore the peptide’s sequence. ^34^ Here, we computed the occupancy of the H-bonds between the Bbp and Fg*α* before the complex rupture. We identified the key amino acid interactions responsible for keeping the complex together at high force loads (Table 1). Different than SdrG:Fg*β*, Bbp:Fg*α* interactions are not dominated by backbone-backbone interactions, with a significant amount of side-chain interaction of the peptide playing an important role in the complex mechanostability. The backbone interactions between Bbp ^*Leu*584, *Thr*582, *Thr*586^ and Fg*α* ^*Thr*565, *Ser*567, *Thr*586^ have been previously described as important for Fg*α* binding at the crystal structure. ^24^ However, we noticed that the side-chain H-bonds are rearranged upon application of mechanical stress on the complex. On the crystal, Bbp^*Asp*334^ forms a side-chain H-bond with Fg*α* ^*Ser*566^, and during the SMD simulations, this interaction shifts to Fg*α* ^*Thr*565^, being the H-bond with the highest occupancy over the trajectories. Another shift occurs between Bbp ^*Asp*334, *Ile*335^ interacting with Fg*α* ^*Phe*564^, on the crystal, to Bbp ^*Ser*333^ interacting with Fg*α* ^*Phe*564^ in our simulations. The H-bond between Bbp ^*Asp*588^ and Fg*α* ^*Gln*563^ is described as important to lock the peptide N-terminus and is still present before the rupture of the complex, although with lower occupancy. Instead, a charged sidechain interaction arises with significant occupancy values: Bbp ^*Asp*556^:Fg*α* ^*Lys*562^. These data corroborates the importance of backbone interactions to maintain the high mechanostability and also highlights important side chain H-bonds plasticity that occurs when Bbp:Fg*α* is exposed to mechanical stress.

**Table 1:** Hydrogen bonds occupancy between Bbp and Fg*α* residues calculated and averaged before the main rupture event.

| Bbp | Fgα | Occupancy (%) | Nature |
| --- | --- | --- | --- |
| Asp334 | Thr565 | 54.84 | Side-chain |
| Asp556 | Lys562 | 45.39 | Side-chain |
| Leu584 | Thr565 | 36.14 | Backbone |
| Thr582 | Ser567 | 35.62 | Backbone |
| Thr586 | Gln563 | 35.03 | Backbone |
| Ser333 | Phe564 | 18.54 | Side-chain |
| Asp588 | Gln563 | 12.77 | Side-chain |
| Thr587 | Ser561 | 12.66 | Side-chain |

### The force propagates indirectly from the latch to the peptide

How a shear force load “activates” the hyperstability of the complex can be investigated by analysing the evolution of pairwise interactions during the force-loading event. Such analysis can be used to investigate how a catch-bond may be formed in the Bbp:Fg*α* complex. ^42^ Previously, it has been shown that SdrG:Fg*β* presents a catch-bond behavior,^41^ which is expected also for Bbp:Fg*α*. To analyse the pairwise interactions during the SMD, we employed the generalized correlation-based dynamical network analysis method, ^56^ which can also be used to calculate force propagation pathways.^44^ Figure 4A shows the pairwise interactions obtained from the network analysis. The thickness of the connections between nodes (amino acid residues) represents how well correlated the motion of these nodes are, and therefore how well connected are these amino acid residues.

**Figure 4:**
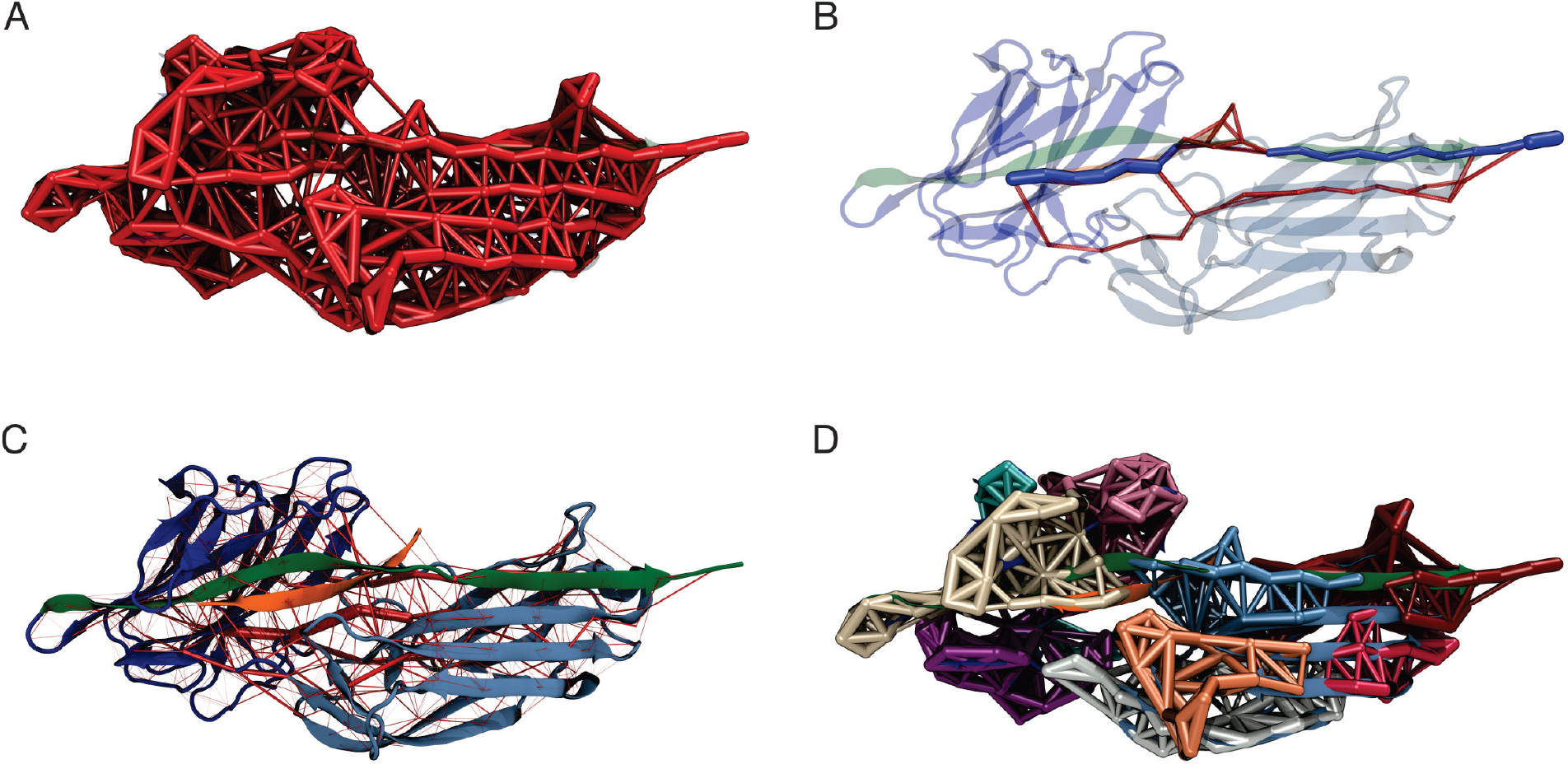
Bbp:Fg*α* dynamical network under high mechanical load. **a)** Representation of the dynamical network. The thickness of the links between the nodes (amino acid residues) represents the correlation of motion between these residues. **b)** The force propagates from the latch indirectly to the peptide, passing by the N2 domain of the protein. The color scheme of the complex is the same from Figure 1. The network’s optimal path is colored in dark blue while the sub-optimal paths are colored in red. **c)** Full dynamical network revealing the most correlated regions of the complex. The weight of the network edges (represented by the thickness of red tubes) is given by the betweenness values. **d)** Generalized correlation-based communities represented by different colors of the nodes and edges in the network.

The force propagation pathway that connects the pulling and the anchoring residues shows that most of the force is propagating from the protein latch directly to the peptide, passing by the center of Bbp’s N2 domain (Figure 4B). These results are slightly different than the ones obtained for the SdrG:Fg*β* complex upon high mechanical stress.^34^ However, in a previous study, it was observed that changes in the pulling velocities can lead to different force propagation pathways, suggesting different unbinding mechanisms at different pulling rates.^40^

The rigidity of the protein under high-force load can also be studied using the betweenness map from the dynamical network analysis (see Figure 4C). The betweenness is defined as the number of shortest paths from all vertices to all others that pass through that node, in this case, an amino acid residue. If an amino acid residue has high betweenness, it tends to be important for controlling inter-domain communication within a protein. ^56^ High betweenness values (thicker red tubes) are seen on the latch that is in direct contact with Fg*α*, highlighting the strong correlation between the motif and the peptide. Interestingly, high betweenness is also found at connections intra N2 domain, pointing that Bbp:Fg*α* complex becomes more rigid under high force loads, particularly in the region interconnecting the latch, the peptide and the N2 domains. Such behavior helps the stabilization of the interactions under high forces.

A representation of the network in subgroups, or communities, is shown at Figure 4D. The communities group the amino acid residues that are most inter-connected in relation to the rest of the network. We can see that Bbp:Fg*α* is subdivided in a handful of communities. The latch, most of Fg*α*, and part of the N2 domains are united in the same community in light blue, showing that these amino acids are highly connected. We also measured the correlation between motions on the interface residues to determine how cooperative their motion is and the essential contacts that are keeping the complex stable under high mechanical load. Essentially, the higher the correlation between residues, the more relevant is their interaction for the stability of the protein complex. We noticed that two Fg*α* residues are highly correlated (values equal or superior to 0.5) to Bbp at the interface, namely: Fg*α* ^*Gln*563^: Bbp ^*Asp*588, *Ser*585, *T hr*586, *T hr*587^ and Fg*α* ^*Phe*564^:Bbp ^*Ser*585^ (Figure 5). The importance of Fg*α* ^*Gln*563^ described as a persistent H-bond contact with Bbp ^*Asp*588, *Thr*586^ and important locking contact is reinforced by its high correlation values.

**Figure 5:**
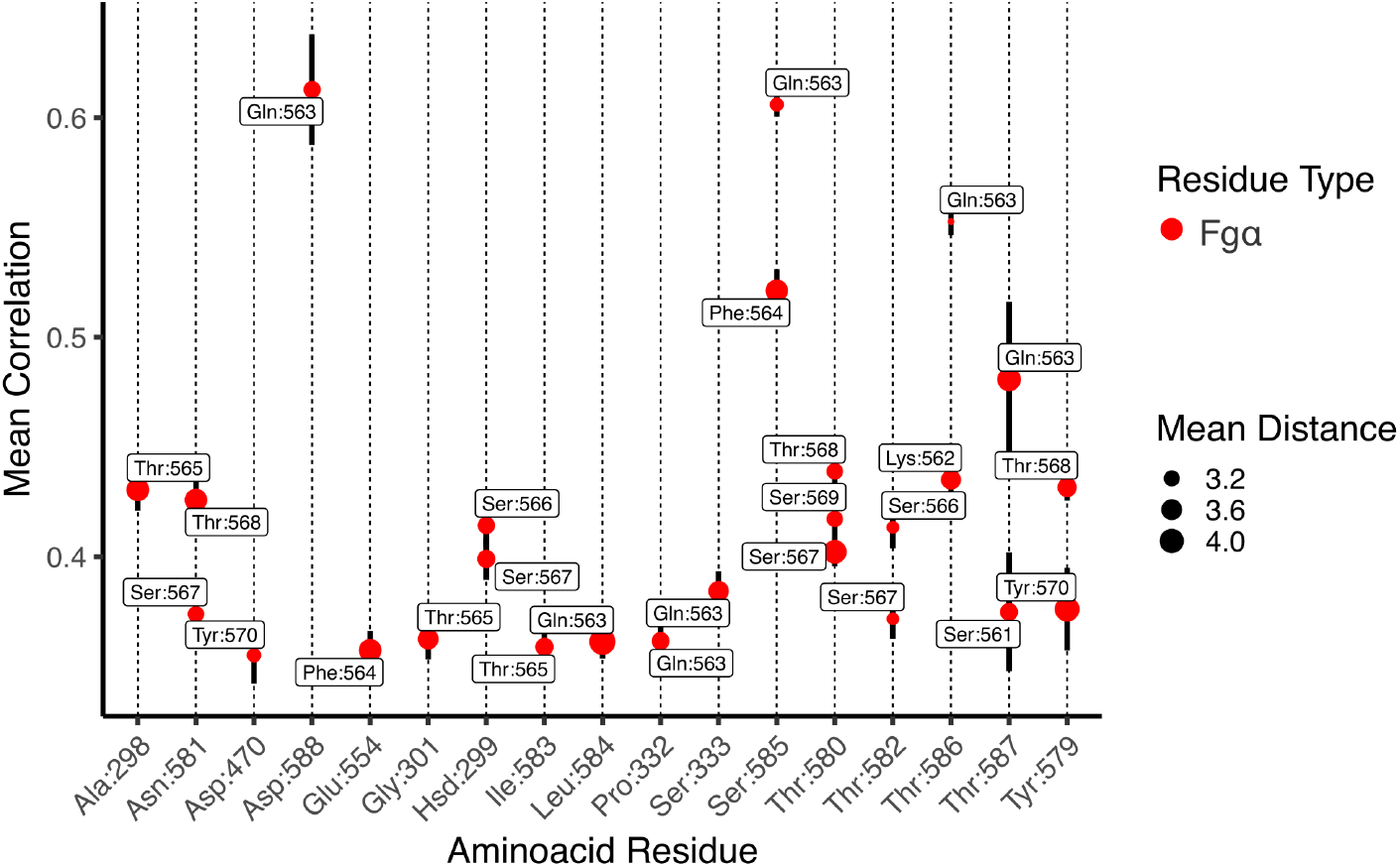
Mean generalized coefficients for contacts along Bbp:Fg*α* inerface. The *x* axis is labeled by Bbp amino acid residues and the *y* axis indicated the averaged generalized correlation values (vertical bars indicate the standard error of the mean), labeled by Fg*α* aminoacid residues. The circle sizes indicates the average Cartesian distance. Only amino acid residues with a mean correlation higher than 0.35 are shown.

## Discussion

During infection, Gram-positive bacteria are frequently exposed to high mechanical stress. These bacteria have evolved an intricate host-binding mechanism to efficiently form colonies under the most inhospitable conditions. Key for the maintenance of the colonies, biofilms are an important virulence factor developed by *S. aureus* among other bacteria. In the initial steps of infection and biofilm formation, MSCRAMMS adhesins have an important role in clinging the bacteria to their human hosts.^7,16^ *S. aureus* isolated from patients suffering from septic arthritis and osteomyelitis specifically interacts with bone sialoprotein, present at bone and dentine extracellular matrix. This interaction is mediated by an specific adhesin protein, namely Bbp.^23,57,58^ Here we have explored the interaction of Bbp with Fg*α* by using an *in silico* SMFS approach that relies on aa- and CG-SMD simulations. CG-SMD simulations have proven to bridge the force-loading gap between *in vitro* SMFS data with *in silico* data obtained from aa-SMD simulations, distanced by orders of magnitude.^54^ In addition, CG-SMD simulations require much less computational power, ^59,60^ enabling us to explore pulling speeds unfeasible to simulate via aa-SMD.^54^

Here, we showed that Bbp:Fg*α* complex can withstand forces even higher than the previously investigated SdrG:Fg*β* complex. ^34^ We revealed that the force propagation pathway between the anchoring and pulling points of the Bbp:Fg*α* complex goes beyond the interactions between the latch and the peptide, passing through an intricate network involving several amino acids of the Bbp N2 domain (Figure 4). We were also able to point the key residues H-bonds responsible for keeping the complex stable at such high mechanical stress, highlighting important backbone-backbone interactions between Bbp ^*Leu*584, *Thr*582, *Thr*586^ and Fg*α* ^*Thr*565, *Ser*567, *Thr*586^ but also side-chain connections, such as Bbp ^*Asp*334^:Fg*α* ^*Thr*565^, Bbp ^*Ser*333^:Fg*α* ^*Phe*564^ and Bbp ^*Asp*588^:Fg*α* ^*Gln*563^ (Table 1). The latter being an important contact to lock the peptide N-terminus. ^24^ Fg*α* ^*Gln*563^ has also revealed to be a key network hub, being highly correlated with several residues on the complex interface such as Bbp ^*Asp*588, *Ser*585, *T hr*586, *T hr*587^ (Figure 5).

In summary, by probing the Bbp:Fg*α* complex under high mechanical load, we discovered the molecular mechanism that triggers Bbp’s unique resilience to shear forces. The high force-loads that can be found during initial stages of bacterial infection stabilize the interconnection between the protein’s amino acids, particularly along the *β*-sheets that, due to their force-loading geometry, cannot be “peeled” like other *β*-sheet-rich proteins, such as green fluorescent protein (GFP)^61,62^ and human filamins.^38,39^ Our results build on previous knowledge of host-microbial interactions, supporting the idea that anti-adhesion therapies might be fundamental in our fight against nosocomial bacteria infections. The detailed molecular picture provided here is critical for the intelligent design of peptidomimetic drugs that can offer the same intramolecular pairwise connections, a key step in creating a new class of antibiotics that act on the initial stages of bacterial infection.

## Methods

### All-atom molecular dynamics simulations

The structure of Bbp in complex with Fg*α* has been previously solved by means of X-ray crystallography at 1.45 Å resolution.^24^ Here we retrieved this structure from the Protein Data Bank (PDB ID: 5CFA) and prepared it for molecular dynamics (MD) simulations using using VMD^63^ and its plugin QwikMD.^64^ The complex was solvated using the TIP3 water model,^65^ with the net charge of the protein neutralized using a 150 mM concentration of sodium chloride.

Steered molecular dynamics (SMD) simulations were carried out using NAMD 3,^66^ with the CHARMM36 force field.^67^ The simulations were performed assuming periodic boundary conditions in the isothermal-isobaric ensemble (NPT) with temperature maintained at 300 K using Langevin dynamics for temperature and pressure coupling, the latter kept at 1 bar. A distance cut-off of 11.0 Å was applied to short-range non-bonded interactions, whereas long-range electrostatic interactions were treated using the particle-mesh Ewald (PME)^68^ method. Taking advantage of a hydrogen-mass repartitioning method implemented in VMD’s autopsfgen, the time step of integration was chosen to be 4 fs for all production aa-MD simulations performed. Before the SMD simulations, the system was submitted to an energy minimization protocol for 1,000 steps. An MD simulation with position restraints in the protein backbone atoms was performed for 1 ns, with temperature ramping from 0 K to 300 K in the first 0.5 ns at a timestep of 2.0 fs in the NVT ensemble, which served to pre-equilibrate the system. In an *in silico* single molecule force spectroscopy (SMFS) strategy, ^43,69^ SMD simulations were carried out in several replicas, using a constant velocity stretching protocol at three different pulling speeds of (Table 2). SMD was employed by harmonically restraining the position of the amino acid at the C-ter of Bbp and moving a second restraint point at the C-ter of Fg*α* peptide with a 5 kcal/mol Å^2^ spring constant, with constant velocity in the z axis. The force applied to the harmonic spring is then monitored during the time of the SMD. The pulling point was moved with constant velocity along the z-axis and due to the single anchoring point and the single pulling point the system is quickly aligned along the z-axis. The number of replicas for each velocity is indicated at Table 2.

**Table 2:** Pulling speeds and number of replicas used for the different MD protocols.

| Protocol | Pulling speed (nm/ps) | Number of replicas |
| --- | --- | --- |
| aa-SMD | $2.5 \times 10^{-01}$ | 192 |
| aa-SMD | $2.5 \times 10^{-02}$ | 192 |
| aa-SMD | $2.5 \times 10^{-03}$ | 160 |
| aa-SMD | $1.25 \times 10^{-03}$ | 160 |
| aa-SMD | $2.5 \times 10^{-04}$ | 160 |
| CG-SMD | $2.5 \times 10^{-03}$ | 128 |
| CG-SMD | $2.5 \times 10^{-04}$ | 128 |
| CG-SMD | $2.5 \times 10^{-05}$ | 64 |
| CG-SMD | $2.5 \times 10^{-06}$ | 64 |

### Coarse-grained molecular dynamics simulations

The atomistic model of Bbp:Fg*α* was modeled onto the Martini 3.0 Coarse-grained (CG) force field (v.3.0.b.3.2)^70^ using martinize2 v0.7.3.^71^ A set of native contacts, based on the rCSU+OV contact map protocol, was computed from the equilibrated all-atom structure using the rCSU server^72^ and used to determine Gö-MARTINI interactions^73^ used to restraint the secondary and tertiary structures with the effective depth *ϵ* of Lennard-Jones potential set to 9.414 kJ.mol^−1^. All CG-MD simulations were performed using GROMACS version 2021.5.^74^ The Bbp:Fg*α* complex was centered in a rectangular box measuring with 10.0, 10.0, nm to the x,y, and z directions. The anchor (Bbp C-terminal) and pulling (peptide C-terminal) backbone (BB) atoms were used to align the protein to the Z axis. The box was then solvated with Martini3 water molecules. Systems were minimized for 10,000 steps with steepest descent, followed by a 10 ns equilibration at the NPT ensemble using the Berendsen thermostat at 298K, while pressure was kept at 1 bar with compressibility set to 3e^−4^bar^−1^, using the Berendsen barostat. A time step of 10 fs was used to integrate the equations of motion. Pulling simulations were subsequently done at the NVT ensemble with a time step of 20 fs. The temperature was controlled using the v-rescale thermostat ^75^ with a coupling time of 1 ps. For all CG-MD simulations, the cutoff distance for Coulombic and Lennard-Jones interactions was set to 1.1 nm,^76^ with the long-range Coulomb interactions treated by a reaction field (RF)^77^ with *ϵ*_*r*_=15. The Verlet neighbor search^78^ was used in combination with the neighbor list, updated every 20 steps. The LINCS^79^ algorithm was used to constrain the bonds and the leapfrog integration algorithm for the solution of the equations of motion. Several replicas of CG-SMD simulations were performed at a range of speeds described at Table 2.

### Simulation data analysis

Hydrogen bonds occupancy were calculated and averaged for aa-MD simulations 1 ns before the main rupture event, using VMD.^63^ Mean correlation and dynamical network pathways were calculated using the generalized dynamical network analysis^56^ and VMD for aa-SMD at pulling velocity of 2.5 *×* 10^−4^ nm/ps. In this analysis, a network is defined as a set of nodes that represent amino acid residues, and the node’s position is mapped to that of the residue’s *α*-carbon. Edges connect pairs of nodes if their corresponding residues are in contact, and 2 non-consecutive residues are said to be in contact if they are within 4.5 Å of each other for at least 75% of analyzed frames. The interface residues between Bbp:Fg*α* were defined in a radius of 10 Å between nodes in each molecule. A representative for the full-network, optimal and suboptimal paths and communities was rendered using one of the SMD trajectory replicas. The mean correlation analysis was carried out 1ns before the first rupture event using a cutoff of 0.35 for the mean correlation coefficients. All charts were generated using in-house python scripts. The protein image was rendered using VMD.

## Conflict of Interest Statement

The authors declare that the research was conducted in the absence of any commercial or financial relationships that could be construed as a potential conflict of interest.

## Author Contributions

PSFCG, MF and MP contributed to performing simulations, analysing data, and writing of the manuscript. DG contributed to analysing data and discussion. RB contributed to writing and discussion on *in silico* force spectroscopy, proof-reading, manuscript revision and approval of the submitted version. PSFCG and RB coordinated the project.

## Funding

This work was supported by the National Science Foundation under Grant MCB-2143787 (CAREER: *In Silico* Single-Molecule Force Spectroscopy) and ACCESS Allocations (Project: BIO220009). We thank Auburn University and the College of Sciences and Mathematics for the computational resources provided by Dr. Bernardi faculty startup funds.

## Acknowledgments

We thank Dr. Marcelo Melo for the fruitful discussions.

## Data Availability Statement

The datasets for this study can be available upon reasonable request to the corresponding author.

## Notes

### Competing Interest Statement

The authors have declared no competing interest.

